# Multi-omic integrated curvature study on pan-cancer genomic data

**DOI:** 10.1101/2022.03.24.485712

**Authors:** Jiening Zhu, Anh Phong Tran, Joseph O. Deasy, Allen Tannenbaum

## Abstract

In this work, we introduce a new mathematical framework based on network curvature to extract significant cancer subtypes from multi-omics data. This extends our previous work that was based on analyzing a fixed single-omics data class (e,g, CNA, gene expression, etc.). Notably, we are able to show that this new methodology provided us with significant survival differences on Kaplan-Meier curves across almost every cancer that we considered. Moreover, the variances in Ollivier-Ricci curvature was explored to investigate its usefulness in network topology analysis as this curvature may be capturing subtle functional changes between various cancer subtypes.

## 1 Introduction

The development of next-generation gene sequencing technologies has allowed researchers to create large multi-omics datasets such as the Cancer Genome Atlas (TCGA), which contain a wide array of information on copy number alteration, DNA methylation, and mRNA expression of a large number of human genes [13]. Uncovering how these cancers alter the normal functioning of key regulatory pathways in the human body is an important step in finding new treatment targets. Within each cancer type, there are cancer subtypes that may lead to dramatic differences in how patients respond to various cancer therapies. The classification and understanding of these subtypes remains incomplete, remains a major challenge in cancer research.

In our previous work, we introduced the use of curvature, a mathematical concept originating from Riemannian geometry [9], to cluster cancer subtypes in an unsupervised way. Our past work seems to suggest that there is a positive correlation between curvature and the robustness of regulatory gene network, and these curvature values can be used to readily classify samples within a cancer type into significantly distinct clusters. In addition to our application of curvature to cluster various cancerous datasets, the concept of curvature also finds its way in numerous applications such as computer vision, signal processing, and machine learning [2, 4, 20]. Anish et al. used curvature to measure changes in robustness of brain networks in children with autism spectrum disorder [16]. Elkin et al. found correlation between Ricci curvature and distinct survival outcomes for a set of patients with ovarian cancer using protein-protein interactive (PPI) networks and gene copy number alterations (CNAs) [7].

In the present paper, we explore a certain extension of Ricci curvature on graphs to study the robustness of combined multi-omics data. By constructing a multi-layer graph combining CNA, methylation, and RNA (gene expression), we study the combined robustness properties of cancer networks. As a feature describing robustness, we cluster pan cancer patients into subgroups based on the computed curvature derived from cancer PPI networks, which yields significant survival differences across a wide range of common cancers. In the following sections, we introduce our methodology then give the application to cancer employing data from TCGA. Finally, we conclude with some future directions of research.

## 2 Background and Methods

### 2.1 Curvature

Curvature is a basic mathematical concept which describes the shape of a manifold by quantifying its deviation from being Euclidean. We refer the interested reader to the standard text [6, 9] for a detailed discussion of Riemannian curvature, and the derived notions of sectional, Ricci, and scalar curvature. In addition to its theoretical significance, curvature is of great importance in numerous applications such as computer vision, signal processing and machine learning [2, 4, 20]. Ricci curvature is also strongly related to entropy and therefore information theory via the Wasserstein distance on the space of probablity distributions on a Riemannian manifold [11]. In some seminal work, Yann Ollivier found a natural notion of Ricci curvature to discrete graphs based on optimal transport, which is widely used in network theory including community detection and feature extraction [15]. In the following subsections, we will briefly review the concept of optimal transport and Wasserstein distance, and then give the formal definitions of Ollivier-Ricci curvature.

### 2.2 Wasserstein distance

The French civil engineer and mathematician, Gaspard Monge first formulated the problem of optimal mass transport in 1781[18, 19]. It was inspired by the need to find the optimal path (relative to a given cost functional) for moving a pile of soil from a given point to a final one in a mass preserving manner. Leonid Kantorovich [10] later reformulated and relaxed the Monge formulation, reducing it to one of linear programming problem. One may show that Kantorovich and Monge formulations are equivalent in a number of cases under certain continuity constraints; see [18, 19] and the references therein. We will follow the Kantorovich formulation in this paper.

In the discrete case, the problem may be defined as follows:

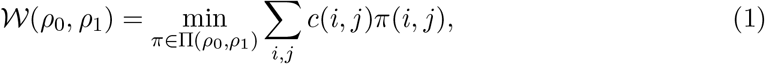

where Π(*ρ*_0_, *ρ*_1_) is the set of matrices 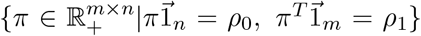, and 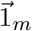 and 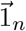 are vectors all of whose entries are of length *m* and *n*, respectively. *c*(*i*, *j*) is the cost of moving a unit mass from location *i* in the original space to location *j* in the target space. The objective function describes exactly the total cost of moving the initial distribution(*ρ*_0_) to the target distribution(*ρ*_1_). Note that *c*(·, ·) can be any nonnegative convex function. In particular, if *c* is chosen to be the distance of the given metric space, then the transport cost is called the 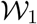 Wasserstein distance or the Earth Mover’s Distance (EMD).

### 2.3 Ollivier-Ricci Curvature

Consider a weighted graph 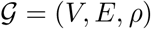. A Markov chain process may be defined as follows, for each vertex *i*, there is an associated probability distribution *μ_i_* given by:

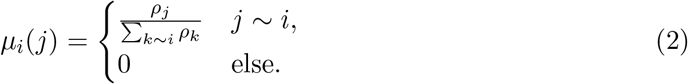

The *Ollivier-Ricci curvature* for *i*, *j* ∈ *V* is defined as:

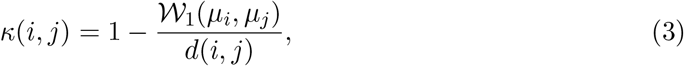

where 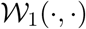 is the Wasserstein distance on the graph and *d*(*i*, *j*) is the shortest distance on the graph between vertices *i* and *j*.

Ricci curvature characterizes the average distance of geodesic balls in a Riemannian manifold. In a flat space, the latter distance is the distance from the centers *d*, while in the positive curvature or negative curvature cases one gets a measure smaller than or greater than *d*, respectively [9]. As noted in [17], this may be expressed in terms of balls equipped with a uniform measure on the the given manifold from which (3) is inspired. On weighted graphs, positive curvature is connected with the number of invariant triangles, while negative curvature with bridges [1]. See Figure 1.

**Figure 1:**
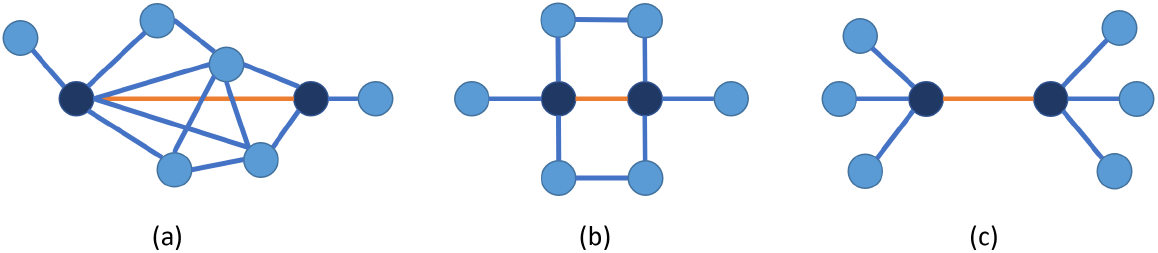
Examples of Ollivier-Ricci curvature of orange edges: (a) positive, neighboring nodes are connected, hub-like (b) zero (c) negative, neighboring nodes are not connected, tree-like

### 2.4 Multi-layered graphs and multi-omics integrated curvature

#### 2.4.1 Single-omic case

There have been a number of works in which the Ollivier-Ricci curvature is applied to single-omic cancer networks expressed as weighted graphs (equivalently Markov chains under certain constraints).

Considering each gene as a vertex on the graph and each edge as a direct interaction between connected genes, we use the PPI network from Human Protein Reference Database (HPRD) [14]. The edges are given by all pairs of genes that have a known protein-protein interaction. With the network topology fixed for each sample (patient), the omics data is then used to calculate the edge weights. This process can be repeated for any given omic modality: copy number alteration (CNA), DNA methylation, and mRNA gene expression. For cases in which there are negative node weight values, one may exponentiate or translate to get all positive weights.

In addition to the nodal weights, edge weights are derived from the nodal values in the following manner: for an edge (*i*, *j*) ∈ *E*, the edge weight *w_ij_* is

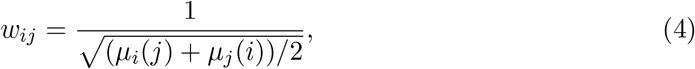

where *μ_i_*(*j*) is defined in (2). These edge weights are used as intrinsic distances on the graph for the 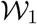 calculation. Specifically, *d*(*i*, *j*) is taken to be the length of the shortest weighted path on the graph, computed efficiently by using the dynamic linear programming based Floyd-Warshall algorithm, to recursively find the shortest distances between all pairs. [3].

With all above definitions, the Ollivier-Ricci curvature may be computed for each edge as specified in equation (3).

#### 2.4.2 Multi-omics case

When more than one type of genomic data is given, we want to employ an appropriate version of Ollivier-Ricci curvature to study the given network in an integrated manner.

Suppose that for each gene *i*, we have a vector-valued weight, 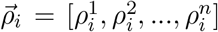, where 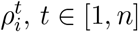 may specify a given nodal feature (e.g., CNA, mRNA, methylation). We want compute the curvature along all the edges in *E* integrating the genomic features utilizing the PPI network as well as the relationships among the features. We propose to accomplish this by imposing a Cartesian product graph structure. Given to two graphs 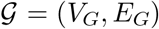 and 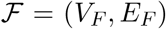, the Cartesian product graph 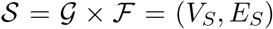 is a graph whose nodes are Cartesian product of the nodes from 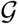 and 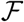:

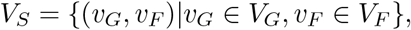

and its edges are defined as:

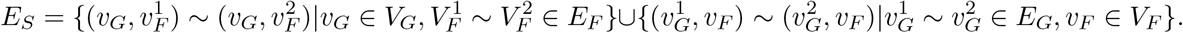

Here 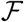 is a graph connecting the layers (equal to *n* here) and 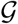 is a PPI network as above.

We give an example for *n* = 3 in Fig 2. Here we illustrate the very important biological case where CNA, DNA methylation and mRNA gene expression are integrated to form a three-layer Cartesian product graph, where the CNA layer directly connects to the mRNA layer, and the methylation layer directly connects to mRNA layer. Note that the CNA and methylation layers are not directly connected, since at present it is not known if CNA and methylation have a direct effect on one another.

**Figure 2:**
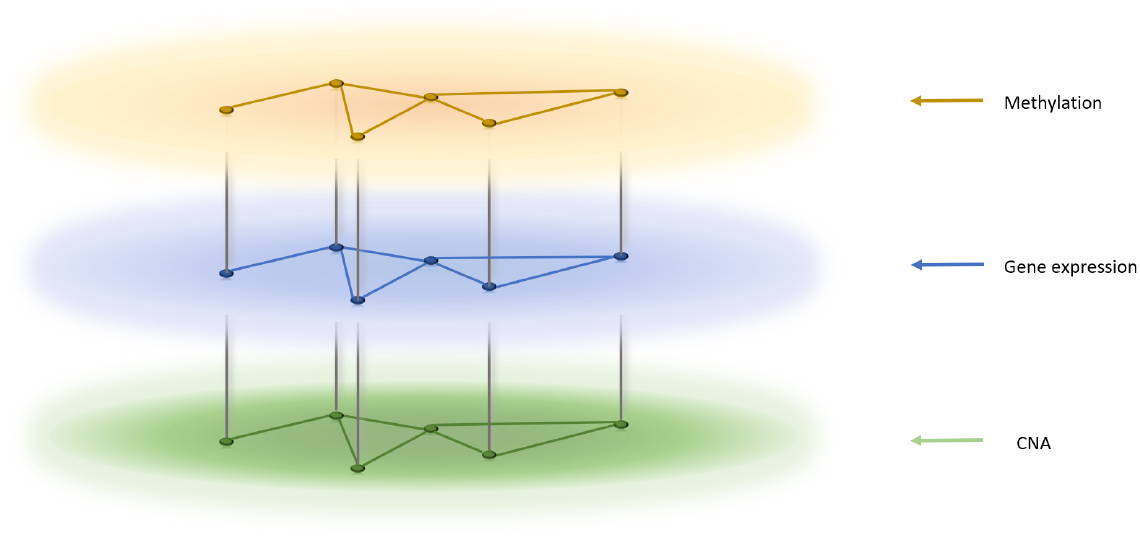
A three-layer Cartesian product graph

Following the same procedure as the one-layer case, we first define a random walk 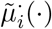 for the *n* layers:

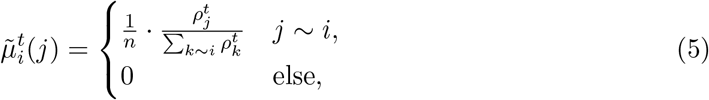

where 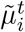 is the part of distribution that is located in layer *t* ∈ [1, *n*].

Next, we define edge weights for the Cartesian product graph. There are two types of edges:

1. Edges that lie within each layer:

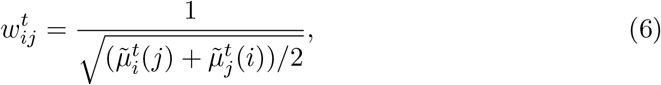
2. Edges that connect different layers:

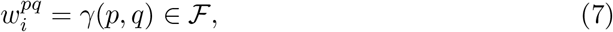

where *γ* is a constant that is fixed for all the inter-layer edges, and may be used to control the strength of the inter-layer interaction.

With all the edge weights defined above, the transport problem on the Cartesian product graph is computed giving the multi-omics integrated extension of Ollivier-Ricci curvature:

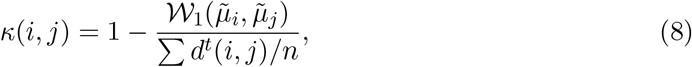

where *d^t^*(*i*, *j*) is taken to be the length of weighted shortest path of the *t*-th layer of the Cartesian product graph.

**Figure 3:**
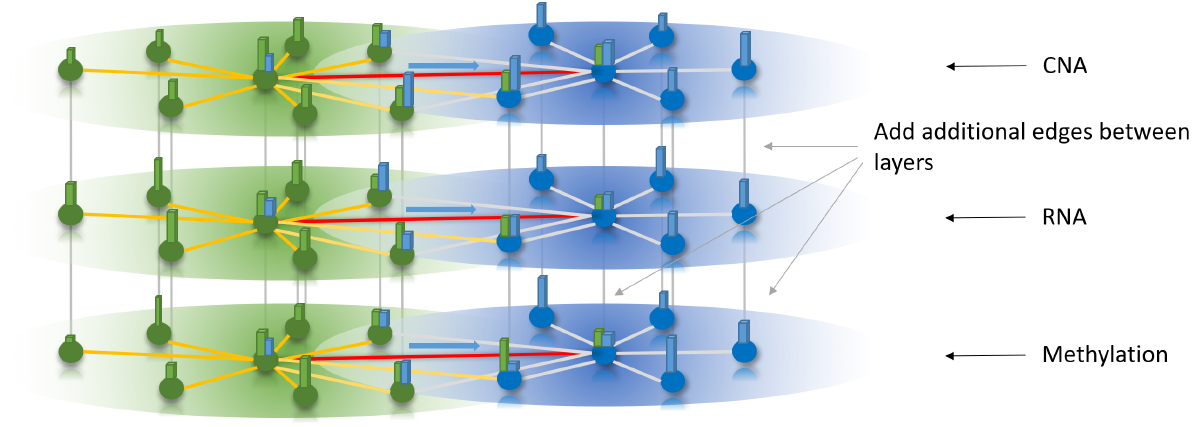
An illustration of vector based curvature

### 2.5 Variants of Ollivier-Ricci Curvature

Despite utility of the original Ollivier-Ricci curvature as well as our extension to the multi-layer Cartesian product graph case, we found some variants of the original formulation very useful. The first is the random walk with residue, which is widely used in many previous studies [8]. For the second variant, we extend the idea of “comparing to Euclidean” notion of curvature to “comparing to any base space.”

#### 2.5.1 Random walk with residue

The random walk in (2) and (5) may be modified as follows:

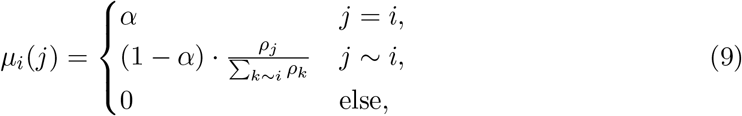

and similarly,

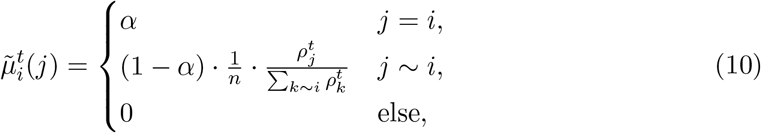

where *α* ∈ [0, 1] is a constant. When *α* = 0, (9) and (10) are the same as (2) and (5). This may be understood as a “half-step” random walk in the original definition, which is a common practice in many applications.

#### 2.5.2 Deviation from a baseline space other than Euclidean

We first consider once again the formula of Ollivier-Ricci curvature (3). The numerator is the actual cost of moving the probability density at a given node to another while the denominator is the intrinsic distance on the graph (there are several possibilities).

We propose to define a version of Ollivier-Ricci curvature as follows:

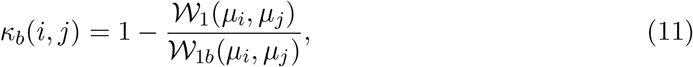

where 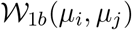 is the 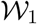 transport cost of moving the neighborhood of *i* to neighborhood of *j* in a fixed baseline space. For this work, we chose

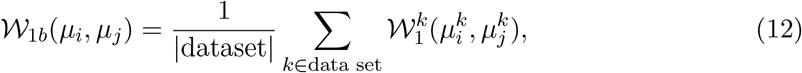

the average value of all the samples in a data set. In the context of cancer, researchers are often concerned about how a regulatory pathway deviates from a normal healthy state. The mean value of all samples in a data set give us an estimate of the base space or normal state. The idea is to make those normal behavior gene interactions to have near zero curvature. Now curvature measures a space that is more hub-like or bridge-like than the given baseline space.

Notice that in this variant we only linearly move the zero point of curvature. With this change, those edges with drastic change in robustness among all patients in the dataset are transformed to very positive or negative.

## 3 Results

### 3.1 Kaplan-Meier results

We applied the above methodology on TCGA datasets for several common major cancers. The multi-omics data were downloaded from the cBioPortal database and provided information about CNA, RNA-seq gene expression, and hm450 DNA methylation data. We also employed the Metabric breast cancer data in our study. Here we only had microarray gene expression and CNA. For all the datasets, The interaction network was derived from the HPRD PPI network [14]. For each cancer dataset, we first found the genes that are common to all available omics data. We then took the intersection of the resulting gene list with that of HPRD, and finally studied the largest connected component of the graph. The numbers of genes and samples for all processed data sets are listed below.

**Table 1:** Basic information of data sets we use

| Dataset name | Cancer | # of layers | # of genes | # of samples |
| --- | --- | --- | --- | --- |
| TCGA_brca | Breast Invasive Carcinoma | 3 | 2992 | 712 |
| Metabric | Breast Invasive Carcinoma | 2 | 3147 | 1904 |
| TCGA_hnsc | Head and Neck Squamous Cell Carcinoma | 3 | 3157 | 388 |
| TCGA_paad | Pancreatic Adenocarcinoma | 3 | 3187 | 137 |
| TCGA_luad | Lung Adenocarcinoma | 3 | 3087 | 376 |
| TCGA_laml | Acute Myeloid Leukemia | 3 | 2864 | 155 |
| TCGA_ov | Ovarian Serous Cystadenocarcinoma | 3 | 2956 | 237 |
| TCGA_gbm | Glioblastoma multiforme | 3 | 2840 | 204 |
| TCGA_skcm | Skin Cutaneous Melanoma | 3 | 3090 | 315 |

By combining the curvature results with a standard hierarchical clustering technique [12], we generated clusters with significant differences in survival as shown through the Kaplan-Meier estimator across practically all cancers on which we tested our methodology. This indicates that the network curvature may contain important information regarding subtype differences within a given cancer type. These clusters can then be used to investigate possible biological differences exist between various cancers.

Gene regulatory networks capture complicated biology. The change of one gene might be compensated by other genes. So analyzing genes or each gene type individually might not yield satisfactory results. Instead, our curvature analysis considers the connections between genes via their neighbors. The values of curvature give a measure of local geometry characterizing the subtle difference among the samples. By clustering patients into subgroups in terms of curvature, patients in the same cluster tend to have similar local geometry, which is a more fundamental local feature. The differences of curvature may correspond to gene functional changes, which reflects survival differences among subgroups.

**Figure 4:**
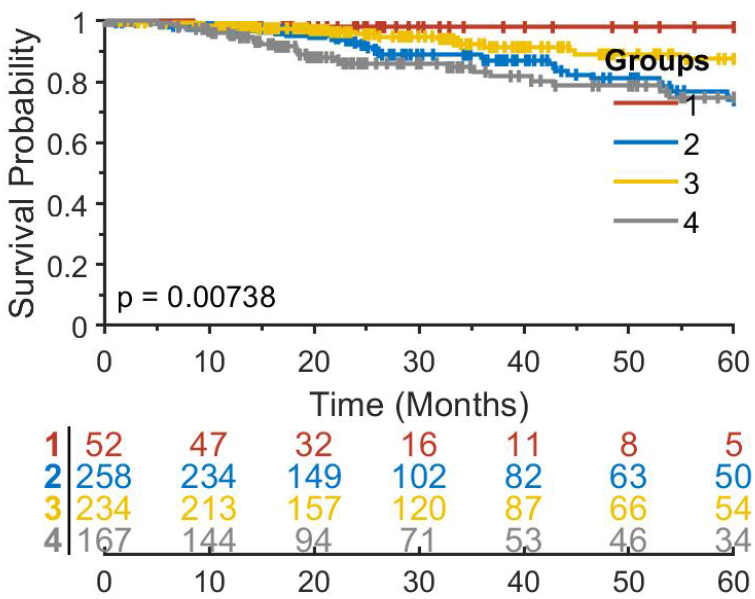
Kaplan-Meier results for TCGA breast cancer data set with *α* = 0.5, *γ* = 1.

**Figure 5:**
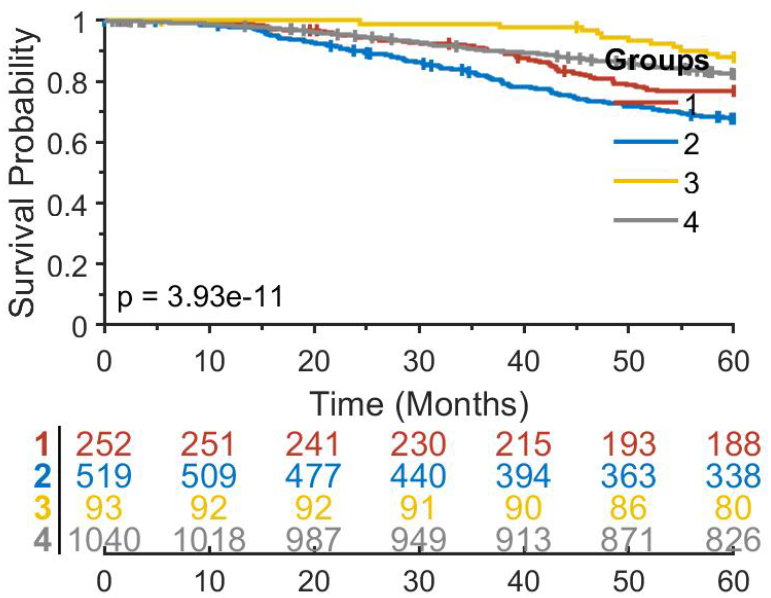
Kaplan-Meier results for Metabric breast cancer data set with *α* = 0.5, *γ* = 1.

**Figure 6:**
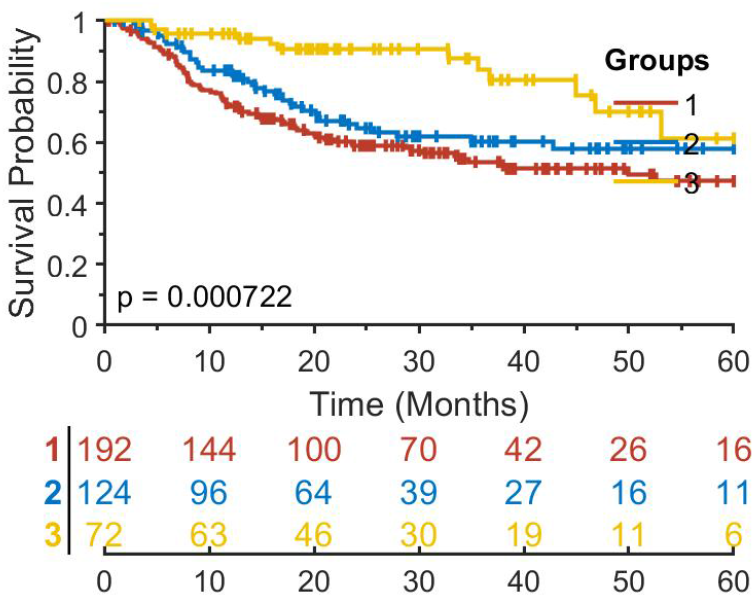
Kaplan-Meier results for TCGA head and neck cancer data set with *α* = 0, *γ* = 1.5.

**Figure 7:**
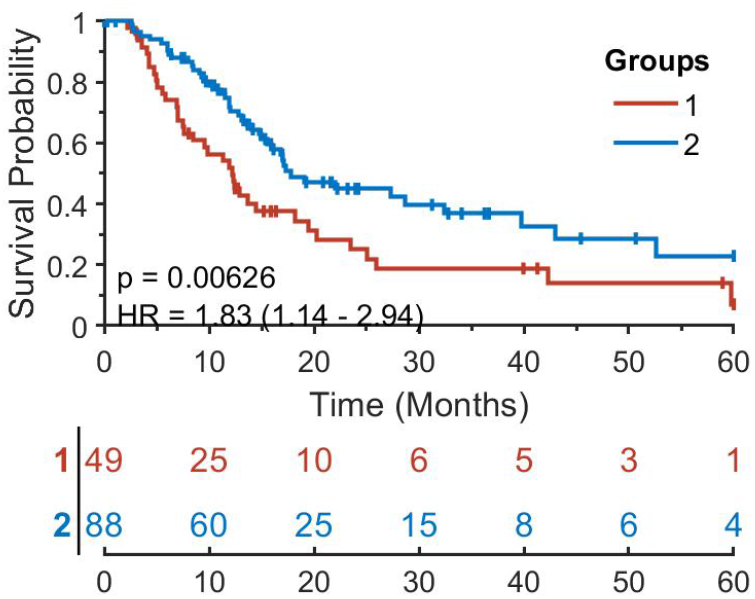
Kaplan-Meier results for TCGA pancreatic cancer data set with *α* = 0.5, *γ* = 1.

**Figure 8:**
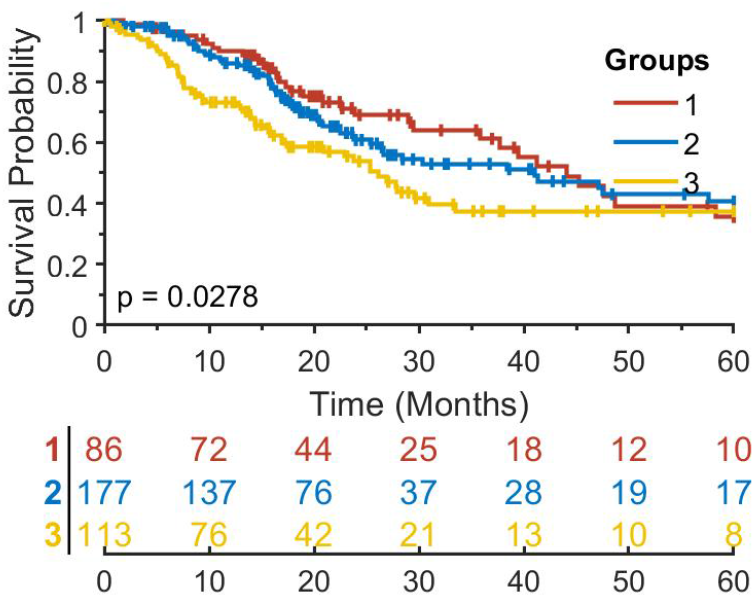
Kaplan-Meier results for TCGA lung adenocarcinoma data set with *α* = 0.5, *γ* = 1.

**Figure 9:**
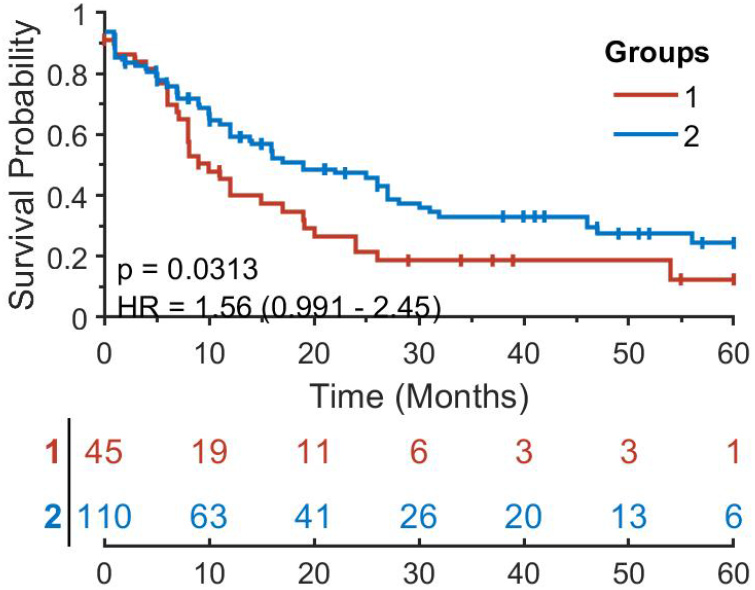
Kaplan-Meier results for TCGA leukemia data set with *α* = 0, *γ* = 0.1.

**Figure 10:**
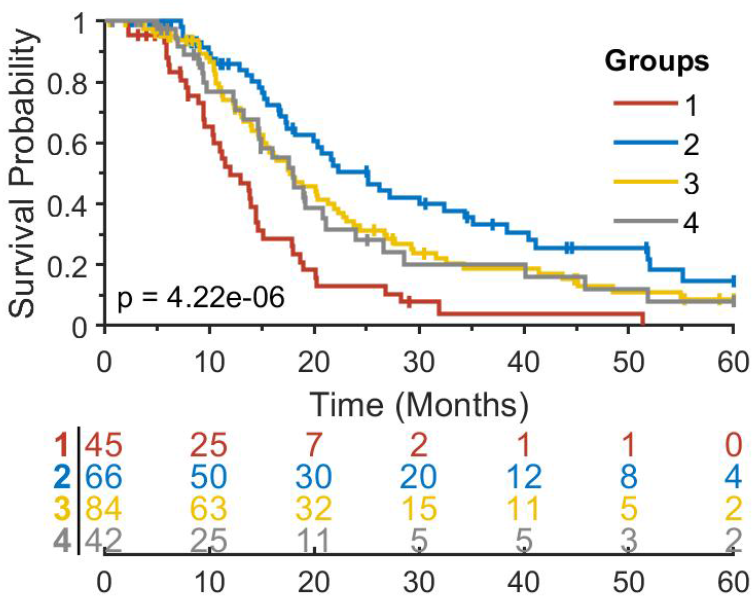
Kaplan-Meier results for TCGA ovarian cancer data set with *α* = 0, *γ* = 1.5.

**Figure 11:**
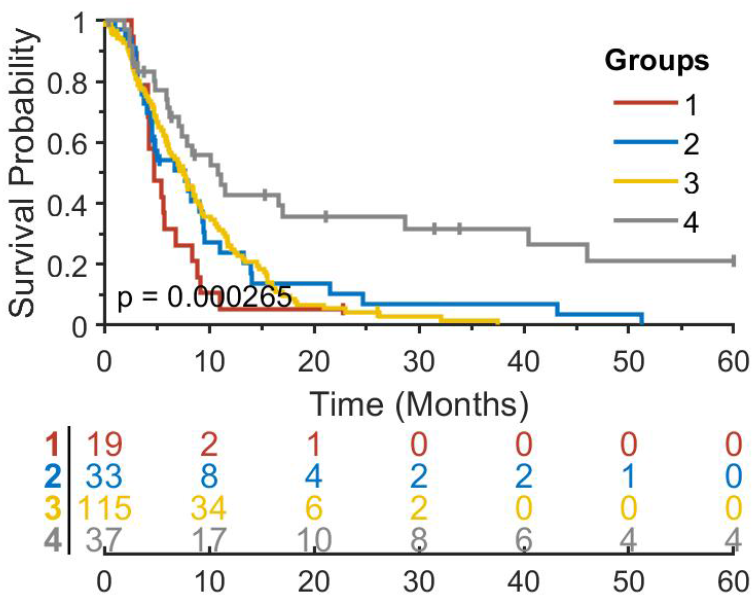
Kaplan-Meier results for TCGA glioblastoma data set with *α* = 0.25, *γ* = 2.

**Figure 12:**
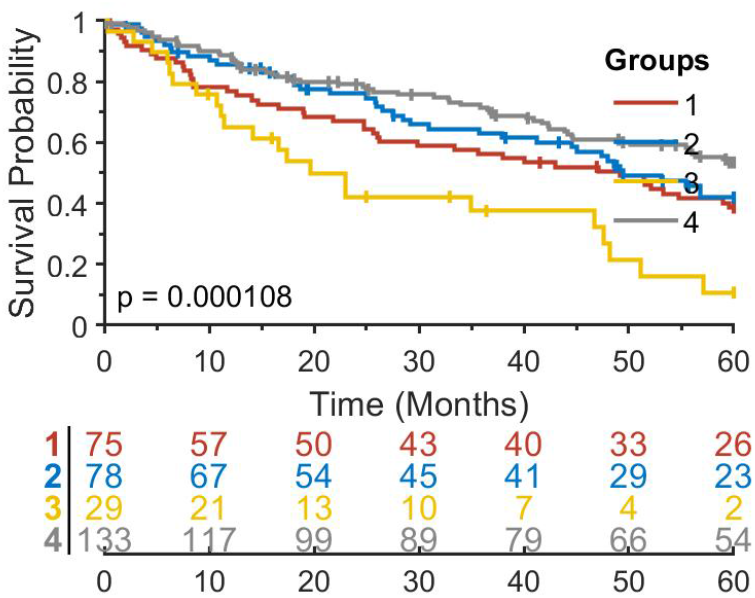
Kaplan-Meier results for TCGA skin cutaneous melanoma data set with *α* = 0, *γ* = 1.5.

## 3.2 Significant curvature difference in different subtypes of cancer

For several well-known cancer subtypes, we investigated whether our curvature method could capture biological differences from the network perspective. Focusing on the breast cancer Metabric dataset, the most malignant subtype of breast cancer is regarded to be triple negative breast cancer meaning that all three ER, PR and Her2 statuses are negative. Most of the patients with this subtype have mutant TP53, while the majority of breast cancer patients that have an ER/PR positive subtype have wild-type TP53. We found that, in terms of curvature values (11), TP53 switched from very negative in the triple negative subtype to very positive in ER/PR positive set.

Mutations of TP53 may change its functions. Indeed, TP53 may lose its ability to suppress tumor growth for many cancer types [5]. We found that the curvature values in different subtypes of breast cancer are very different, sometimes quite dramatic.

**Table 2:** Top 20 negative genes in Metabric triple negative, TP53 mutant subtype

| Top genes | Curvature |
| --- | --- |
| CREBBP | -5.41835 |
| UBE2I | -2.40985 |
| SMAD3 | -1.91895 |
| UNC119 | -1.72264 |
| BRCA1 | -1.40683 |
| SRC | -1.20088 |
| TP53 | -1.09668 |
| RARA | -1.08374 |
| CALM1 | -1.03648 |
| USP7 | -1.02801 |
| YWHAB | -0.99047 |
| CALCOCO2 | -0.9844 |
| AKT1 | -0.96324 |
| CDK2 | -0.93956 |
| ERBB2 | -0.92471 |
| PLK1 | -0.92258 |
| PRKCB | -0.91013 |
| PTPN11 | -0.86984 |
| GNB5 | -0.86932 |
| RB1 | -0.81533 |

**Table 3:** Top 20 positive genes in Metabric triple negative, TP53 mutant subtype

| Top genes | Curvature |
| --- | --- |
| VIM | 2.607904 |
| PTPN6 | 1.887768 |
| PTK2 | 1.749174 |
| ATN1 | 1.669488 |
| JUN | 1.527905 |
| JAK2 | 1.393305 |
| EP300 | 1.343139 |
| CSNK2A2 | 1.278762 |
| ITGB1 | 0.947019 |
| RNF11 | 0.871667 |
| TNFRSF1A | 0.856385 |
| APP | 0.831585 |
| CDK1 | 0.806475 |
| JAK1 | 0.802274 |
| BCAR1 | 0.793185 |
| PRKACA | 0.786608 |
| SVIL | 0.78544 |
| ZHX1 | 0.751178 |
| XRCC6 | 0.734371 |
| IRS2 | 0.647708 |

**Table 4:** Top20 negative genes in Metabric ER/PR positive, TP53 mutant subtype

| Top genes | Curvature |
| --- | --- |
| PTK2 | -1.04547 |
| SRC | -0.77467 |
| CSNK2A2 | -0.70023 |
| EP300 | -0.65179 |
| YWHAB | -0.63889 |
| VIM | -0.59992 |
| PRKCA | -0.56983 |
| ERBB2 | -0.474 |
| BCAR1 | -0.45246 |
| PLCG1 | -0.43309 |
| ZHX1 | -0.42126 |
| XRCC6 | -0.38781 |
| GRB2 | -0.37808 |
| ITGB2 | -0.3732 |
| SIAH1 | -0.31029 |
| PICK1 | -0.30819 |
| MMP2 | -0.30534 |
| CDK1 | -0.28164 |
| PTPN6 | -0.28121 |
| APP | -0.27914 |

**Table 5:** Top20 positive genes in Metabric ER/PR positive, TP53 mutant subtype

| Top genes | Curvature |
| --- | --- |
| CREBBP | 3.02489 |
| TP53 | 2.071297 |
| UBE2I | 1.30434 |
| BRCA1 | 0.88627 |
| PLK1 | 0.708575 |
| DLG4 | 0.698465 |
| SMAD4 | 0.693955 |
| SMAD2 | 0.606423 |
| SMAD3 | 0.587212 |
| PRKCB | 0.529638 |
| USP7 | 0.524551 |
| BCL2 | 0.490532 |
| STAT3 | 0.477343 |
| POLR2A | 0.463449 |
| PDPK1 | 0.398286 |
| RB1 | 0.38917 |
| FXR2 | 0.360223 |
| CALM1 | 0.33756 |
| HRAS | 0.333579 |
| KPNB1 | 0.328164 |

## 4 Discussion and future work

As a mathematical concept, curvature has been studied extensively and applied to various problems in biology and medicine, in particular cancer. In the present paper, we proposed an extension of Ollivier-Ricci curvature which integrates more than one type of omics data. The proposed curvature represents the robustness in terms of multi-level genomic data. Experiments indicate that curvature differences may be be strongly indicative of survival differences. Further, the change of curvature sign of key genes may indicate change of function from wild type to mutant in different subtypes of cancers. In our paper, we only integrated two or three types of genomic data, which are most readily accessible. Other data types such as mutation status may also be included in our framework, and will be the topic of future work.

We studied curvature for each edge of a network. In principle, curvature can be computed between any two genes in a network. To study longer range interactions might recover some new information. The success of the application of the geometric notion of network curvature to cancer indicates its relationship to gene functionality and network functional robustness. We plan to extend this work to include the integration of networks at various scales including imagery as well as pathology.

## 5 Acknowledgments

This study was supported by AFOSR grant (FA9550-20-1-0029), NIH grant (R01-AG048769), MSK Cancer Center Support Grant/Core Grant (P30 CA008748), and a grant from Breast Cancer Research Foundation (grant BCRF-17-193).

